# Elucidating Microbial Iron Corrosion Mechanisms with a Hydrogenase-Deficient Strain of *Desulfovibrio vulgaris*

**DOI:** 10.1101/2024.03.24.586472

**Authors:** Di Wang, Toshiyuki Ueki, Peiyu Ma, Dake Xu, Derek R. Lovley

## Abstract

Sulfate-reducing microorganisms extensively contribute to the corrosion of ferrous metal infrastructure. There is substantial debate over their corrosion mechanisms. We investigated Fe^0^ corrosion with *Desulfovibrio vulgaris*, the sulfate reducer most often employed in corrosion studies. Cultures were grown with both lactate and Fe^0^ as potential electron donors to replicate the common environmental condition in which organic substrates help fuel the growth of corrosive microbes. Fe^0^ was corroded in cultures of a *D. vulgaris* hydrogenase-deficient mutant with the 1:1 correspondence between Fe^0^ loss and H_2_ accumulation expected for Fe^0^ oxidation coupled to H^+^ reduction to H_2_. This result and the extent of sulfate reduction indicated that *D. vulgaris* was not capable of direct Fe^0^-to-microbe electron transfer even though it was provided with a supplementary energy source in the presence of abundant ferrous sulfide. Corrosion in the hydrogenase-deficient mutant cultures was greater than in sterile controls, demonstrating the H_2_ removal was not necessary for the enhanced corrosion observed in the presence of microbes. The parental H_2_-consuming strain corroded more Fe^0^ than the mutant strain, which could be attributed to H_2_ oxidation coupled to sulfate reduction producing sulfide that further stimulated Fe^0^ oxidation. The results suggest that H_2_ consumption is not necessary for microbially enhanced corrosion, but H_2_ oxidation can indirectly promote corrosion by increasing sulfide generation from sulfate reduction. The finding that, *D. vulgaris* was incapable of direct electron uptake from Fe^0^ reaffirms that direct metal-to-microbe electron transfer has yet to be rigorously described in sulfate-reducing microbes.

**Impact Statement:** The economic impact of microbial corrosion of iron-containing metals is substantial. A better understanding of how microbes accelerate corrosion is expected to lead to the development of methods to prevent corrosion. The results presented here refute the suggestion, frequently made in the microbiology literature, that microbial H_2_ uptake stimulates Fe^0^ corrosion. Also refuted, are previous claims that *Desulfovibrio vulgaris* is capable of directly extracting electrons from Fe^0^. The results are consistent with the concept that sulfide produced by sulfate-reducers promotes Fe^0^ oxidation with the production of H_2._ The results illustrate that appropriate mutants can provide rigor to corrosion mechanism studies.

## Introduction

The world-wide economic cost of microbial corrosion is likely to exceed a trillion dollars a year and current strategies for mitigating corrosion are relatively ineffective (1–3). A better understanding of the factors controlling microbial corrosion rates might provide insights for the development of new strategies to prevent corrosion. The microbial impact on the corrosion of ferrous metals is typically most substantial under anaerobic conditions. In anaerobic corrosion Fe^0^ is oxidized to Fe^2+^:

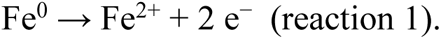

A diversity of microbes can establish electrical contacts with Fe^0^, directly accepting electrons derived from Fe^0^ oxidation to support anaerobic respiration. Microbes capable of this ‘electrobiocorrosion’ (2) include *Geobacter* (4, 5), *Shewanella* (6, 7), *Methanosarcina* (8, 9), *Methanothrix* (9), and *Sporomusa* (9) species. Known electron acceptors for electrobiocorrosion are nitrate, Fe(III), and carbon dioxide. Although the sulfate-reducing microbes *Desulfovibrio ferrophilus* and *Desulfopila corrodens* were proposed to be electrobiocorrosive (10), subsequent studies found that they were not (11).

However, sulfate reducers are often the most abundant and metabolically active microbes associated with intense anaerobic corrosion (12–17). The most abundant sulfate reducers are often *Desulfovibrio* species (18–20). The mechanisms by which *Desulfovibrio* and related sulfate reducers promote corrosion has been a matter of debate since some of the earliest studies of microbial corrosion (19–28). Those early studies often focused on *Desulfovibrio* H_2_ uptake because, even in the absence of microbes, an important route for anaerobic Fe^0^ oxidation is proton reduction:

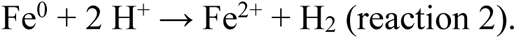

Although it is clear that Fe^0^ oxidation can serve as a H_2_ source for H_2_-oxidizing anaerobes, including *Desulfovibrio* species (see (19, 29) for extensive lists), there has been substantial debate whether microbial H_2_ consumption can accelerate Fe^0^ oxidation coupled to H_2_ production (20–24, 26, 27, 30, 31). Studies conducted under abiotic conditions have indicated that removal of H_2_ should have no effect on the rate of Fe^0^ oxidation with H^+^ reduction and that this reaction is not inhibited as H_2_ concentrations increase (26). Arguments for an important role of microbial H_2_ uptake have often merely demonstrated or inferred that the H_2_ produced from Fe^0^ oxidation was microbially consumed, but did not rigorously demonstrate that H_2_ uptake accelerated Fe^0^ oxidation (20, 22–24).

An example of the difficulty in interpreting the studies on this topic are two studies with the same first author submitted within ten days of each other (24, 32). In the first study submitted, cell suspensions of H_2_-consuming *Desulfovibrio desulfuricans* provided with fumarate as the electron acceptor did not accelerate the weight loss of mild steel over that observed in sterile controls (24). However, the second study reported the results of electrochemical studies conducted with cell suspensions of *Desulfovibrio vulgaris* with benzyl viologen as the electron acceptor that suggested that microbes with hydrogenases could accelerate Fe^0^ oxidation (32).

A complication in evaluating the role of H_2_ uptake in accelerating Fe^0^ oxidation is that the sulfide produced during sulfate reduction can enhance H_2_ production from Fe^0^. Iron sulfides formed from the Fe^2+^ from Fe^0^ oxidation and sulfide, can promote reaction #1 when deposited on Fe^0^ surfaces (27, 28). Furthermore, hydrogen sulfide produced during sulfate reduction can react with Fe^0^ to generate H_2_ (25):

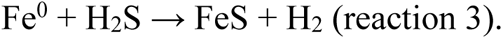

Another potential complicating factor has been the suggestion that iron sulfides might enable direct Fe^0^-to-microbe electron transfer, i.e. electrobiocorrosion (33). This hypothesis was derived from electrochemical studies. Cathodic currents were generated with electrodes poised at (−0.4 V) in the presence of *D. vulgaris* with iron sulfide nanoparticles on the cell surface, but not in the absence of the iron sulfide-coated cells (33). This result was interpreted as direct electrode-to-microbe electron transfer. However, iron sulfides can readily lower electrode overpotentials for the reduction of H^+^ to H_2_ (24, 34) and iron sulfides may stimulate H_2_ uptake by sulfate reducers (34). These potential alternative explanations for the higher cathodic currents in the presence of the iron sulfide-coated cells were not rigorously ruled out. These considerations, and the fact that the role of microbially produced ferrous sulfides on Fe^0^ oxidation was speculated, not experimentally demonstrated, indicates that more rigorous evaluation of the routes for Fe^0^ corrosion in the presence of iron sulfides is required.

An effective approach for evaluating the role of H_2_ uptake in microbial corrosion is to construct strains in which the genes for hydrogenases have been deleted (4–6, 35). For example, studies with strains of *Geobacter sulfurreducens* and *Shewanella oneidensis* in which the genes for the H_2_ uptake hydrogenases were deleted demonstrated that these microbes accepted electrons from Fe^0^ even as H_2_ accumulated (4, 6). However, these microbes were also capable of electrobiocorrosion, providing an alternative route for electron uptake from Fe^0^ (4, 6). In contrast, a hydrogenase-deficient mutant of *D. vulgaris* unable to utilize H_2_ did not reduce sulfate with Fe^0^ as the electron donor whereas the parental strain did (35). This result demonstrated that *D. vulgaris* relied on H_2_ as an intermediary electron carrier between Fe^0^ and cells and was incapable of electrobiocorrosion when only Fe^0^ was available as the electron donor and energy source. However, some methanogens are only effective in electrobiocorrosion when they are metabolizing an additional energy source as they oxidize Fe^0^ (8, 9). The possibility that *D. vulgaris* might also directly accept electrons from Fe^0^ in the presence of an additional energy source was not previously evaluated (35). This is important because microbes also have access to organic substrates in most environments in which metals are being corroded (2).

Here we report on Fe^0^ corrosion studies with the same hydrogenase-deficient strain of *D. vulgaris* employed in previous studies (35), but this time grown with lactate as an additional electron donor. The results demonstrate that microbial consumption of H_2_ consumption was not essential for extensive corrosion during sulfate reduction and that the hydrogenase-deficient strain of *D. vulgaris* was incapable of electrobiocorrosion, even with extensive ferrous sulfide production and the additional lactate energy source. Multiple lines of evidence were consistent with the concept that microbial sulfide production stimulates Fe^0^ oxidation with the production of H_2_.

## Results and Discussion

### Biofilm growth

*D. vulgaris* was grown in the presence of lactate and Fe^0^ as potential energy sources because organic substrates are available to microbes in most anaerobic corrosive environments (2) and because the hydrogenase-deficient mutant does not grow with Fe^0^ as the sole electron donor (35). The parental strain (JW710) and the hydrogenase-deficient mutant strain (JW5095) grow at similar rates in the lactate-sulfate medium employed in this study (35). When Fe^0^ was included in the lactate-sulfate medium, both strains produced substantial biofilms on the Fe^0^ surfaces (Figure 1).

**Figure 1.**
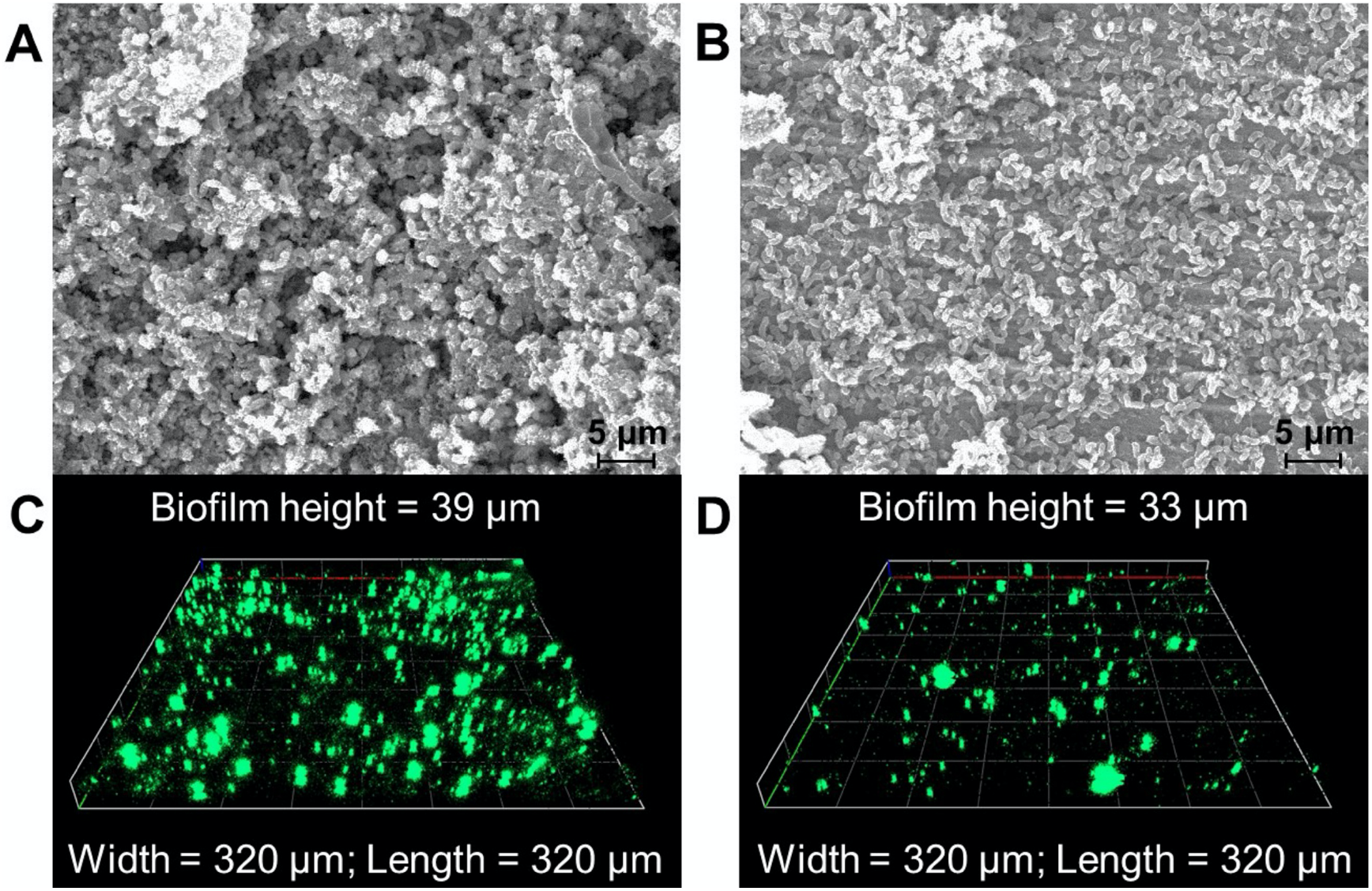
Biofilm growth on Fe^0^ surfaces. Scanning electron microscopy images of the biofilms of the parental (A) and hydrogenase-deficient (B) strains after fixation and gold coating. Confocal laser scanning microscope visualization of parental (C) and hydrogenase-deficient (D) strains after staining with Live/Dead BacLight™ bacterial staining kit.

### Lack of direct electron transfer and hydrogenase impact on corrosion

As expected from previous studies (35), H_2_ accumulated in the hydrogenase-deficient mutant cultures, but not with the parental strain (Figure 2A). The H_2_ produced (Figure 2A) and the Fe^0^ loss (Figure 2B) in the mutant cultures was three-fold higher than that produced in previously reported in sulfide-free sterile incubations under the same conditions (36). This result is consistent with the concept that as microbes reduce sulfate to sulfide, the sulfide promotes Fe^0^ oxidation with the reduction of H^+^ to H_2_. Although some microbes release hydrogenases that can promote Fe^0^ oxidation with the generation of H_2_ (37, 38), this possibility can be ruled out in our studies because the hydrogenase genes had been deleted from the mutant and even the parental strain does not release extracellular hydrogenases (35).

**Figure 2.**
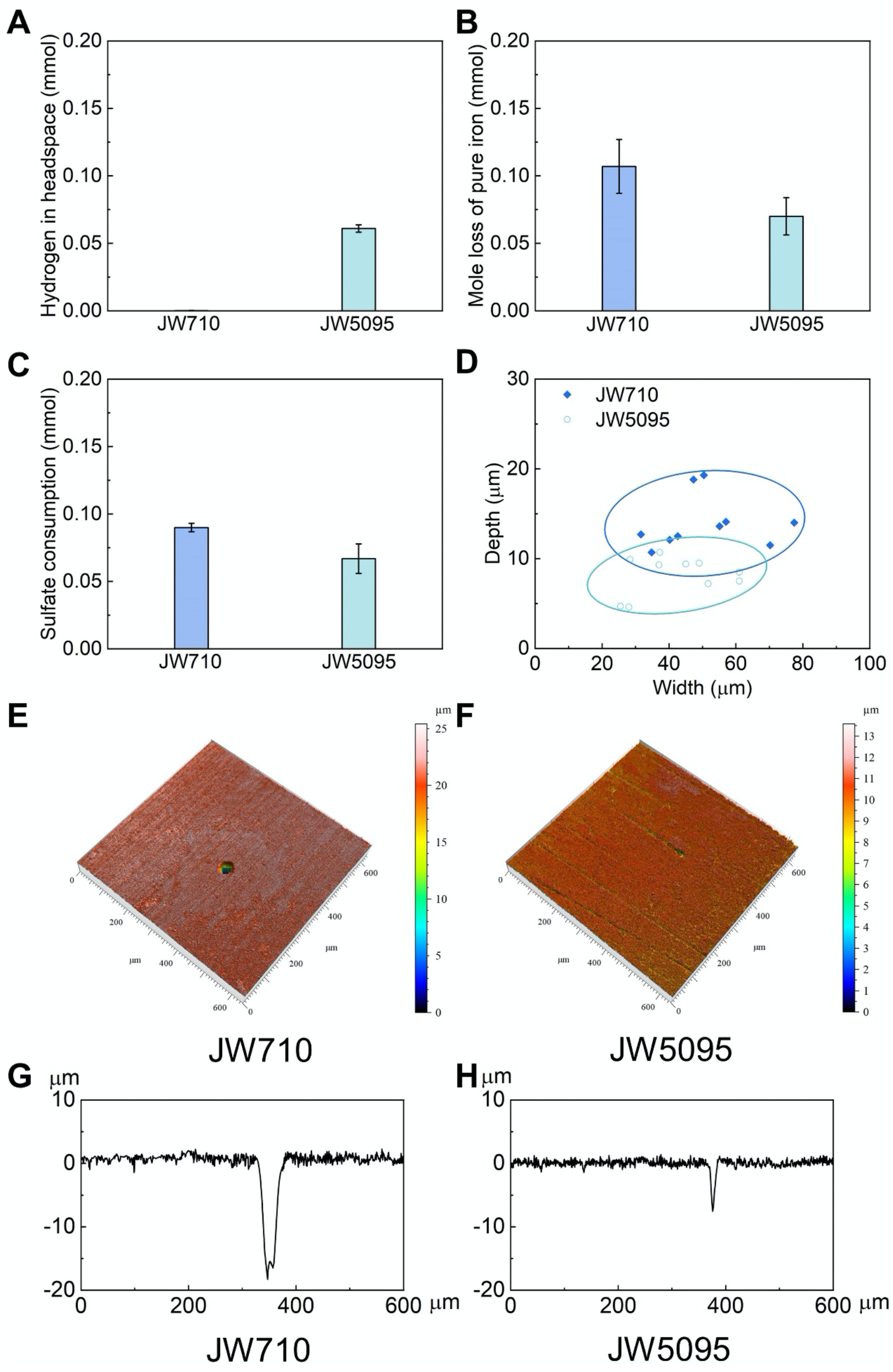
Metabolic and corrosion parameters of the parental strain (JW710) and hydrogenase-deficient strain (JW5095) during growth in lactate-sulfate medium in the presence of Fe^0^ after seven days of incubation. (A) H_2_ accumulation. (B) Loss of Fe^0^. (C) Sulfate consumption. (D) Pit parameters from the ten deepest corrosion pits of the two strains. Surface morphology (E, F) and profiles (G, H) of the deepest pits for each strain. Values in A-C are the mean and standard deviation of triplicate incubations.

Multiple lines of evidence indicated a lack of direct Fe^0^-to-microbe electron transfer. When Fe^0^ is oxidized via direct electron transfer to microbes no H_2_ is produced. However, within the error of the measurements, the accumulation of H_2_ in the hydrogenase-deficient mutant cultures (0.061 ± 0.0028 mmoles (mean ± standard deviation n = 3); Figure 2A) was comparable to the loss of Fe^0^ (0.070 ± 0.012 mmoles; Figure 2B), as expected from the 1:1 correspondence between Fe^0^ oxidation and H_2_ production when Fe^0^ is oxidized with the reduction of H^+^ (reaction #2). Furthermore, if the hydrogenase-deficient strain had derived electrons from direct electron transfer then those electrons would have contributed to sulfate reduction. Yet, when it is considered that ca. 10% of the organic substrate supporting anaerobic respiration is diverted to biosynthesis, the amount of sulfate reduction (0.067 ± 0.011 mmoles of sulfate reduced; Figure 2C) compared well with the 0.0675 mmoles of sulfate expected to be reduced with the 5 mM lactate that was provided as an electron donor (5 mmol lactate/L × 0.9 [fraction consumed in respiration] × 0.03 L of media × 1 mmmol sulfate reduced/2 mmol lactate oxidized to acetate = 0.0675 mmoles sulfate reduced). Thus, the Fe^0^ loss/H_2_ accumulation stoichiometry and the extent of sulfate reduction demonstrate a lack of *D. vulgaris* direct electron uptake even though lactate was available as an additional energy source and abundant ferrous sulfide was produced.

### Dual electron sources for the parental strain

The parental strain reduced more sulfate than the mutant strain (Figure 2C) and substantially more than the sulfate reduction possible with the 0.075 mmol of lactate available. Thus, unlike the mutant strain, the parental strain benefited from electrons derived from Fe^0^ oxidation. The evidence suggests that these electrons were transferred to the parental strain via a H_2_ intermediate. The low H_2_ levels in the parental strain cultures (Figure 2A) demonstrated that the parental strain consumed the H_2_ that accumulated in the mutant cultures. Just the parental strain consumption of that excess H_2_ alone could account for 65 % of the additional sulfate that the parental strain reduced because:

1. The additional sulfate reduction by the parental strain was 0.023 mmol (0.090 mmol sulfate reduced by the parental strain ‒ 0.067 mmol sulfate reduced by the mutant strain = 0.023 mmol)
2. The parental strain consumed the 0.061 mmol of H_2_ that accumulated in the mutant cultures
3. Consumption of 0.061 mmol H_2_ reduces 0.015 mmol sulfate (0.060 mmol H_2_ × mmol sulfate reduced/4 mmol H_2_ oxidized = 0.015 mmol sulfate reduced)
4. 0.015 mmol sulfate reduced = 65% of additional parental strain sulfate reduction (0.015 mmol sulfate reduced from H_2_ consumption/0.023 total sulfate reduced × 100 = 65 %).

Furthermore, the greater sulfide production resulting from the parental strain oxidizing H_2_ could be expected to further stimulate Fe^0^ oxidation with the production of even more H_2_. This was apparent from the greater weight loss of Fe^0^ (Figure 2B) and deeper pitting than in parental strain cultures (Figure 2D-H). The 0.008 mmol of sulfate reduction that could not be accounted for from the consumption of the H_2_ accumulation in the mutant cultures is the equivalent of 0.032 mmol of Fe^0^ oxidized (4 mmol Fe^0^ oxidized per mmol of sulfate reduced). Within the error of the measurements, 0.032 mmol Fe^0^ oxidized is comparable to the difference in the weight loss of the parental (0.107 ± 0.018) and mutant (0.070 ± 0.012) strains. As detailed above, extracellular hydrogenases and direct electron uptake could be ruled out as potential mechanisms for promoting sulfate reduction. Thus, the positive feedback loop of sulfate reduction generating sulfide, resulting in more H_2_ production to support more sulfate reduction is the most likely explanation for the greater corrosion in the parental strain cultures.

### Similar corrosion mechanism in both strains

The open circuit potential during Fe^0^ corrosion was similar for both strains (Figure 3A), as would be expected if Fe^0^ oxidation was proceeding with H_2_ production with both strains. This result also suggested that the accumulation of H_2_ in the hydrogenase mutant did not substantially inhibit Fe^0^ oxidation reactions. Corrosion resistance (Figure 3B) was only slightly higher and corrosion current (Figure 3D) was slightly lower with the hydrogenase-deficient strain, reflecting the somewhat greater corrosion with the parental strain (Figure 2B), which as discussed above, can be most likely attributed to greater sulfide deposition with the parental strain.

**Figure 3.**
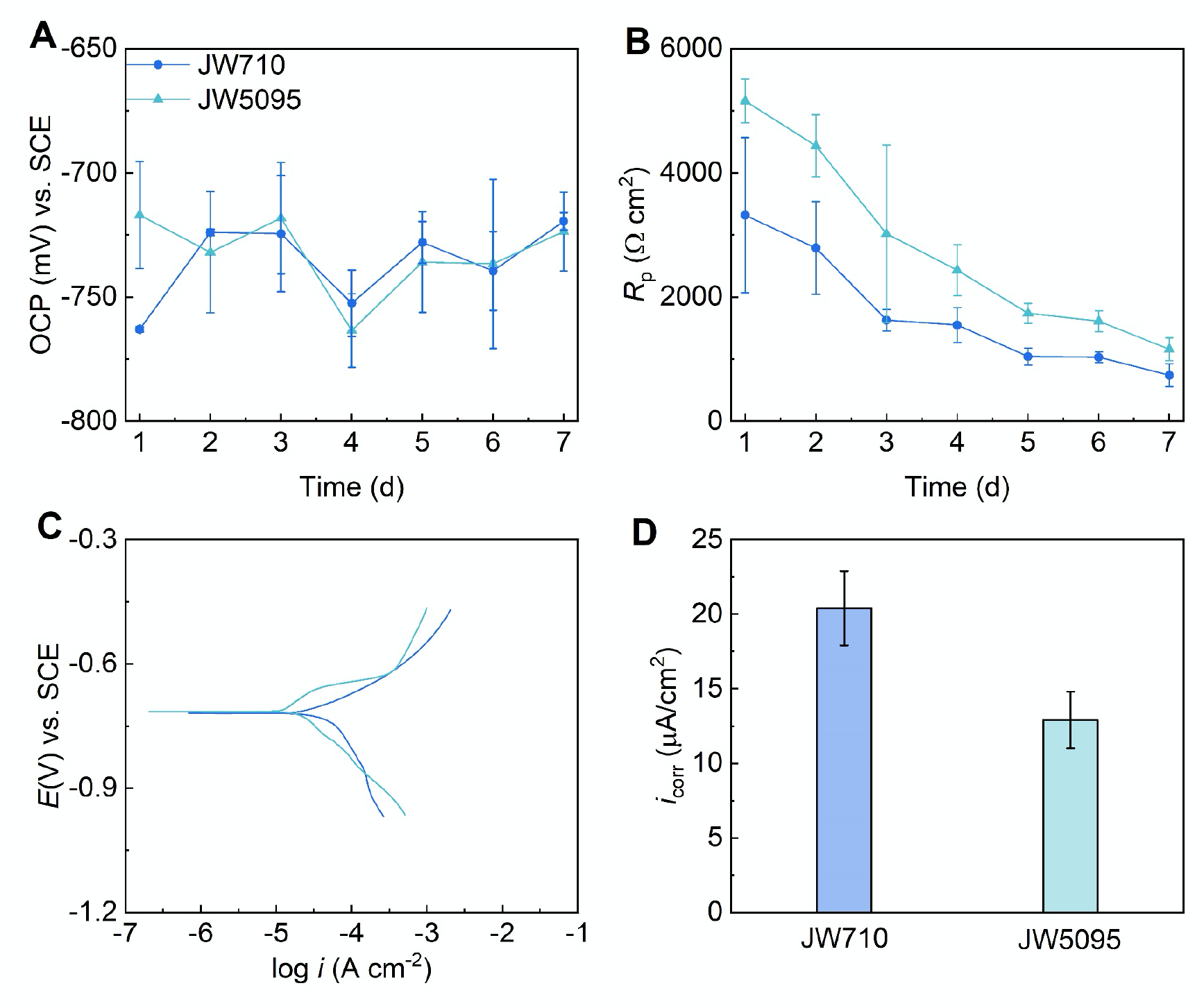
Electrochemical analysis of corrosion with the two strains. (A) Open circuit potential and (B) *R*_p_ during 7-day incubations. (C) Potentiodynamic polarization curves and (D) corrosion current density calculated from potentiodynamic polarization curves on day 7.

## Conclusions

This study illustrates the potential for rigorously evaluating often contentious mechanistic claims in the corrosion literature with mutant strains. The availability of a hydrogenase-deficient mutant enabled studies that clearly demonstrate that *D. vulgaris* is incapable of discernable direct electron uptake from Fe^0^, even when supplied with lactate as an additional energy source and in the presence of abundant ferrous sulfide. The lack of *D. vulgaris* direct electron uptake from Fe^0^ emphasizes that the claim for *D. vulgaris* direct electron uptake from electrodes in the presence of ferrous sulfides (33) should also be reexamined with the hydrogenase-deficient mutant.

The substantial corrosion in cultures of the hydrogenase-deficient mutant provides evidence from a metabolically active system that H_2_ consumption does not promote Fe^0^ oxidation coupled to H^+^ reduction, a conclusion consistent with previous findings in abiotic studies (26). H_2_ partial pressures of more than 2044 atm (207 Mpa), a value much higher than conceivable in any natural environment, would have been required to make Fe^0^ oxidation coupled to H_2_ production (reaction #2) thermodynamically unfavorable in our incubations (See Supplementary Material for calculations).

Thus, H_2_-utilizing sulfate reducers benefit from Fe^0^ corrosion, but do not directly promote corrosion with H_2_ removal. The primary importance of microbial H_2_ consumption is that it provides an electron donor for additional sulfate reduction to sulfide. This was evidenced from the greater corrosion in the presence of the hydrogenase-deficient mutant versus sterile controls, and the higher corrosion in the presence of the parental strain versus the hydrogenase-deficient mutant.

The mechanisms for microbial Fe^0^ corrosion have only been rigorously examined with functional genetic studies in a few model microbes (4–7, 35). Furthermore, other physiological factors such as mechanisms for biofilm formation and how outer-surface cellular components influence Fe^0^-microbe interactions and corrosion have yet to be investigated with in-depth molecular approaches. It is also unknown how well the microbes available in culture represent those that make the largest contributions to Fe^0^ corrosion outside the laboratory (29). A concerted attempt to recover highly corrosive microbes in culture and elucidate how they corrode with definitive genetic and biochemical studies is required if microbial corrosion is to be understood at the level necessary to innovate new corrosion mitigation strategies.

## Materials and Methods

### Strains and growth conditions

Two *D. vulgaris* strains were supplied by Valentine V. Trotter and Adam M. Deutschbauer of the Lawrence Berkeley Laboratory. Strain JW710, also referred to here as the parental strain, is the strain employed for a markerless genetic exchange system in *D. vulgaris* (39). Markerless deletion of all of the known hydrogenase genes in strain JW710 yielded strain JW5095 (40). Previous studies (35) demonstrated that the two strains grow equally well with lactate as the electron donor and sulfate as the electron acceptor. The parental strain JW710 also grows with H_2_ as the sole electron donor whereas JW5095 does not utilize H_2_ (35). Cultures were routinely maintained in the previously described defined NB medium (35) under N_2_/CO_2_ (80:20) with lactate (60 mM) as the electron donor and sulfate (30 mM) as the electron acceptor. Standard anaerobic culture procedures, including maintaining cultures under oxygen-free gases, sealing culture vessels with thick butyl rubber stoppers, and transferring and sampling cultures with needles and syringes were employed throughout.

For corrosion studies the cultures were provided with sulfate (5 mM) as the electron acceptor and lactate (5 mM) and three Fe^0^ coupons (8 mm × 8 mm × 5 mm; wt% composition: Fe 99.95, C 0.01, S 0.008, P 0.01, Si 0.001, Mn 0.001, Ni 0.001, Cr 0.01 and Al 0.006) as potential electron donors in 30 ml of medium in 65 ml serum bottles. Prior to use the Fe^0^ coupons were polished with a sequence of 240, 600, and 1000 grit sandpaper and then washed with ethanol and MilliQ water. The polished coupons were sterilized by soaking them in 75% ethanol and drying them in a UV chamber and then aseptically and anaerobically added to the medium. Incubations were at 30 °C.

### Analytical methods

H_2_ concentrations in the culture headspaces were determined with gas chromatography with a thermal conductivity detector (Trace 1310, Thermo Scientific, USA). For sulfate analysis, liquid samples were diluted in water 20-fold, filtered (0.22 μm pore diameter polyvinylidene difluoride membrane) and analyzed with ion chromatography ((Dionex AS-DV, Thermo Scientific, USA).

### Biofilm visualization

For visualization of biofilms with scanning electron microscopy the Fe^0^ coupons were rinsed with pH 7.4 phosphate buffered solution (PBS) and fixed with glutaraldehyde (2.5% v/v) for 2 h. The samples were then successively dehydrated with 50%, 60%, 70%, 80%, and 90% ethanol solutions for 10 min and then 100% ethanol solution for 5 min before gold-coating the samples and examining them with a EVO 10 scanning electron microscope (Zeiss, Germany).

For biofilm visualization with confocal scanning laser microscopy the Fe^0^ coupons were washed with PBS and then stained with the Live/Dead BacLight™ bacterial staining kit (Molecular Probes L-7012, Thermo Fisher Scientific, USA) for 10 min in the dark. The samples were examined with the fluorescent channels of a LSM900 confocal laser scanning microscope (Zeiss, Germany).

### Analysis of corrosion weight loss and pitting

To determine Fe^0^ weight loss, the coupons were treated with Clarke’s solution (1000 mL hydrochloric acid (specific gravity 1.19), 20 g antinomy trioxide (Sb_2_O_3_) and 50 g stannous chloride (SnCl_2_)) following the ASTM G1-03 protocol (41) to remove corrosion products. The treated surfaces were scanned with the LSM900 microscope in bright field mode and the data analyzed with ConfoMap Premium Software (Zeiss, Germany) to determine corrosion pitting depth, surface morphologies, and pitting profiles.

### Electrochemical characterization

Electrochemical parameters of corrosion were determined with a classical three electrode system with a platinum plate (10 mm × 10 mm × 1 mm) as the counter electrode, a saturated calomel electrode as the reference electrode, and a Fe^0^ electrode with a 1 x 1 cm exposed surface as the working electrode. The open circuit potential (OCP) and linear polarization resistance (LPR) were monitored daily. Potential dynamic polarization (PDP) was determined after seven days of incubation. The electrochemical tests were conducted with electrochemical workstations (Reference 600, Gamry Instruments, USA). The LPR tests were scanned with a rate of 0.167 mV/s in the range of −10 mV to 10 mV versus OCP. PDP were conducted at a scan rate of 0.333 mV/s from −0.3 V to 0.3 V versus OCP.

## Conflict of Interest

The authors declare no conflicts of interest.

## Supplementary Material

### Calculation of the H_2_ Concentration Required For Iron Oxidation Coupled to H_2_ Production to be Thermodynamically Unfavorable

In anaerobic anodic corrosion Fe^0^ is oxidized to Fe^2+^:

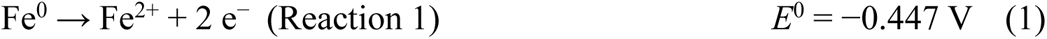

where *E*^0^ is the standard potential.

The proton reduction reaction is:

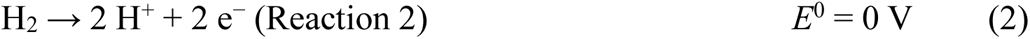

where *E*^0^ is the standard hydrogen electrode at 25 °C, 1.0 M H^+^ and a H_2_ gas pressure of one atmosphere.

The reaction for the oxidation of Fe^0^ coupled to H_2_ production is:

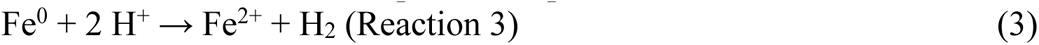

Thus, the potential for Reaction 3 is:

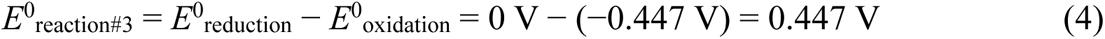

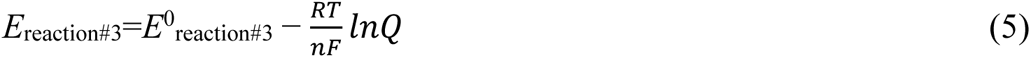

where *R* is the gas constant 8.314 J·K^−1^·mol^−1^, T is the temperature of 30 °C for these studies, *n* is the number of moles of electrons exchanged (2), *F* is Faraday’s constant, 96485 J·V^−1^·mol^−1^, and Q is the reaction quotient of products and reactants.

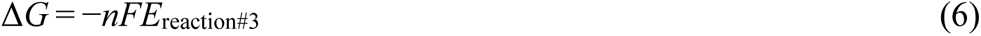

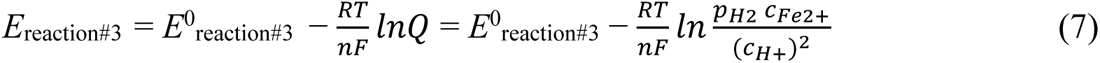

The maximum dissolved Fe^2+^ concentration possible in our incubations would have been generated with the parental strain, which exhibited the most corrosion. The total molar loss of Fe^0^ corroded by parental strain was 0.107 mmol in 30 mL of medium. Most of this the Fe^2+^ generated can be expected to precipitate as iron sulfide, which was abundant in the cultures. However, to be conservative, all the loss of Fe^0^ was considered to be oxidized to dissolved Fe^2+^. Thus, the actual inhibitory H_2_ concentration is expected to be higher than the number calculated here. In the 30 mL of medium, the ideal highest dissolved Fe^2+^ concentration would be 0.107 mmol/0.03mL = 3.57 mM = 0.00357 M. At pH 7 the concentration of H^+^ is 10^−7^ M.

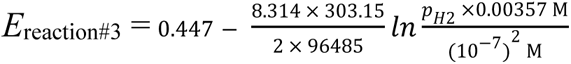

At equilibrium, Δ*G* = 0, and Δ*G* = −*nFE*_reaction#3_, so *E*_reaction#3_ = 0. Thus,

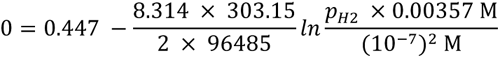

Rearranging:

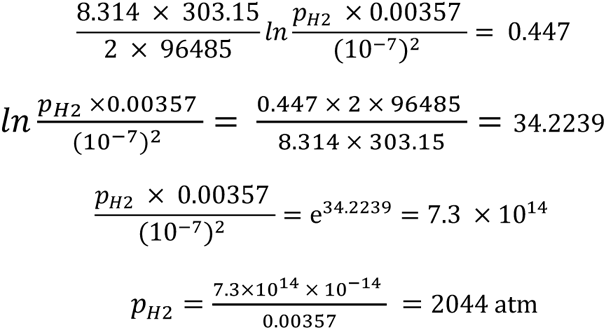

Thus, a conservative estimate is that a H_2_ partial pressure greater than 2044 atm (207 Mpa) would be necessary for Fe^0^ oxidation coupled to H_2_ production to be thermodynamically unfavorable.

## Notes

### Competing Interest Statement

The authors have declared no competing interest.

